# Coevolution does not slow the rate of loss of heterozygosity in a stochastic host-parasite model with constant population size

**DOI:** 10.1101/2020.04.07.024661

**Authors:** Ailene MacPherson, Matthew J. Keeling, Sarah P. Otto

## Abstract

Coevolutionary negative frequency dependent selection has been hypothesized to maintain genetic variation in host and parasites. Despite the extensive literature pertaining to host-parasite coevolution, the effect of matching-alleles (MAM) coevolution on the maintenance of genetic variation has not been explicitly modelled in a finite population. The dynamics of the MAM in an infinite population, in fact, suggests that genetic variation in these coevolving populations behaves neutrally. We find that while this is largely true in finite populations two additional phenomena arise. The first of these effects is that of coevolutionary natural selection on stochastic perturbations in host and pathogen allele frequencies. While this may increase or decrease genetic variation, depending on the parameter conditions, the net effect is small relative to that of the second phenomena. Following fixation in the pathogen, the MAM becomes one of directional selection, which in turn rapidly erodes genetic variation in the host. Hence, rather than maintain it, we find that, on average, matching-alleles coevolution depletes genetic variation.

## 1. Introduction

There is a rich history of evolutionary theory exploring the conditions under which genetic variation is maintained or depleted in finite populations. The loss of genetic variation through drift is exacerbated, for example, by fluctuations in population size (Crow, 1970, eq. 7.6.3.34), but variation is maintained through balancing selection in the form of overdominance (Crow, 1970, eq. 8.6.4) or negative frequency-dependent selection (Takahata and Nei, 1990). First suggested by Haldane (1949), one process that is often posited to maintain genetic variation is coevolution between hosts and their parasites. Coevolution, it is argued, should favour pathogens that are best at infecting the most common host genotype. This in turn should favour the spread of rare host genotypes, a form of negative frequency-dependent selection (NFDS) believed to maintain genetic variation (Clarke, 1979). Despite the long-standing interest in coevolution as a mechanism maintaining variation, few studies have explicitly analysed the dynamics of a host and parasite that are both polymorphic in a fully stochastic framework.

Following Haldane’s initial hypotheses, balancing selection as a result of coevolution in the form of NFDS and/or overdominance was suggested as a mechanism behind the extraordinary genetic diversity found at mammalian Major Histocompatibility Complex (MHC) loci (Bodmer, 1972). These same arguments have been used recently to explain the diversity of anti-microbial peptides in *Drosophila* (Unckless et al., 2016; Chapman et al., 2019). MHC loci, and other immune defence genes, are notable not only for their high levels of heterozygosity (> 200 alleles across three loci Klein and Figueroa, 1986; Zimmer and Emlen, 2013) but also for the long-term trans-specific persistence of these polymorphisms (Lawlor et al., 1988; Klein, 1987). Using a coalescent approach Takahata and Nei (1990) found that heterozygote advantage and NFDS are both capable of generating the observed levels of polymorphism. Importantly, however, the model they used was not explicitly coevolutionary but rather based on NFDS within a single species.

As exemplified by Takahata and Nei (1990), much of the literature on the maintenance of genetic variation alludes to yet blurs the distinction between single-species and coevolutionary NFDS (for example see Tellier et al., 2014; Otto and Michalakis, 1998; Zhao and Waxman, 2016; Llaurens et al., 2017; Ejsmond and Radwan, 2015; Rabajante et al., 2016). By the definition of NFDS in a single-species model (called direct-NFDS, Brown and Tellier, 2011), the fitness of an allele increases as its frequency declines, this in turn can favour the spread of rare alleles and the maintenance of genetic variation. In contrast two-species NFDS (called indirect-NFDS), the sort that commonly arises with host-parasite coevolution, favours alleles of the focal species that correspond to ones which are rare in the interacting species. As noted by Brown and Tellier for the gene-for-gene model (2011), we show that indirect-NFDS in matching-alleles model often has little if any impact on the maintenance of genetic variation.

Hints that indirect-NFDS does not maintain genetic variation can be found throughout the theoretical literature on matching-alleles models. Simulating coevolution in a population where genetic variation is repeatedly introduced through migration or mutation, Frank (1991, 1993) found that the dynamics were dominated by the fixation and loss of genetic variants. He attributed this effect to the repeated population bottlenecks that occur from coevolutionary driven fluctuations in population size, but it was unclear whether this same behaviour would arise in a population of constant size. Similarly, although not explicitly discussed, many individual-based models of coevolution include mutation in either one (Agrawal and Lively, 2002) or both species (Lively, 1999; Borghans et al., 2004; Ejsmond and Radwan, 2015) in order to maintain variation and hence coevolution over the long term. Modelling coevolution in an infinite population, M’Gonigle et al. (2009) found that genetic variation is not maintained at equilibrium except when mutation is very frequent. This is echoed in simulations in finite populations, where in the absence of mutation/migration, fixation in either the host or pathogen is vary rapid (Gokhale et al., 2013; Schenk et al., 2018). In addition to including mutation, there are several theoretical indications that, rather than being driven by NFDS, the emergent effects of coevolution are dependent on the existence of heterozygote advantage in diploids. For example, M’Gonigle and Otto (2011) showed that the evolution of parasitism depends, not solely on NFDS, but on whether the interaction induces heterozygote advantage, on average. Similarly, Nuismer and Otto (2004) showed that whether there is, on average, heterozygote advantage is the key determinant of how ploidy levels evolve in both hosts and parasites.

Despite this long and extensive history of verbal and theoretical models, it is unclear whether indirect-NFDS can indeed maintain genetic variation and explain the extensive heterozygosity observed at immune defence loci. Here we compare the maintenance of genetic variation in finite coevolving populations relative to that expected under neutral drift. Previous studies of finite coevolving populations have focused instead on the number of alleles maintained by mutation and coevolution (Borghans et al., 2004), the time until loss of alleles (Gokhale et al., 2013), and the impact of different model assumptions on the time until loss (Schenk et al., 2018). A direct comparison with the neutral expectation is, however, only possible analytically in simple models of host-parasite coevolution. We aim to understand the effect of host-parasite interactions on the maintenance of genetic diversity by examining one such simple single-locus model of coevolution.

## 2. Theoretical Background

There are two classic models of coevolution involving a single locus major-effect genes in each species, the gene-for-gene (GFG) model, which was motivated by the genetic architecture of flax-rust interactions (Flor, 1956), and the matching-alleles model (MAM), a form of host-parasite specificity that may arise from lock and key molecular interactions (Dybdahl et al., 2014). While GFG and MAM represent only two of the many possible models of host-parasite specificity, their dynamics are representative of the full range that can arise in single-locus models (Agrawal and Lively, 2002; Engelstädter, 2015). In the GFG model, hosts carry either a “susceptible” or “resistant” allele. Susceptible hosts can be infected by both “virulent” and “avirulent” parasite genotypes whereas resistant hosts can only be infected by the virulent parasite genotype. This asymmetry favours fixation of resistant hosts followed by the fixation of virulent pathogens. Deterministic models have shown that this coevolutionary arms race does not in general maintain genetic variation (Leonard, 1977, 1994). Rather stability of genetic polymorphism and the long-term maintenance of genetic variation requires the introduction of temporal delays and asynchrony between host and parasite life cycles (Brown and Tellier, 2011). As GFG is often used to model coevolution in plant-pathogen systems, this asynchrony can be introduced, for example, through seed dormancy or perenniality (Tellier and Brown, 2009).

In contrast to the coevolutionary arms-race dynamics with GFG the symmetry of MAM generates Red Queen allele frequency cycles (also known as trench-warfare dynamics), which are more amenable to the maintenance of genetic variation. MAM is often formulated, and simulated, as an interaction between a host and pathogen with discrete non-overlapping generations (Nuismer, 2017). In each generation hosts are exposed to a single parasite. If host and parasite carry the same, “matching”, allele at the coevolutionary locus then infection occurs with probability *X* whereas interactions between “mis-matching” host and parasite genotypes occur at a reduced rate *Y*. Successful infection decreases host fitness by *α*_*H*_ and increases parasite fitness by *α*_*P*_. The host and pathogen population sizes are often assumed to remain constant in these models such that hosts and pathogens reproduce proportionally to their fitness creating a subsequent generation of the same size.

In the deterministic (infinite population size) limit, the resulting recursion equations for the MAM are characterized by an unstable cyclic equilibrium about a host and pathogen allele frequency of 1*/*2 as shown in Figure 1A (M’Gonigle et al. (2009): haploid host, haploid pathogen, Leonard (1994): diploid host, haploid pathogen). The corresponding dynamics of host heterozygosity is shown in Figure 1B. Starting from a small perturbation near the polymorphic equilibrium, heterozygosity in the host population begins near its maximum of 0.5 and declines, on average, as the allele frequencies cycle outward. As noted above, although NFDS generates Red Queen allele frequency cycles in the short term, these cycles grow in amplitude over time and are not expected to maintain genetic variation in the long term except in the presence of very high mutation rates (M’Gonigle et al., 2009). Indeed, simulating the discrete-time MAM in a finite population, we confirm that host heterozygosity in this model, defined as *H* = 2*p*(1 − *p*) where *p* is the host allele frequency, declines faster than expected under neutral genetic drift (see supplementary *Mathematica* notebook).

**Figure 1:**
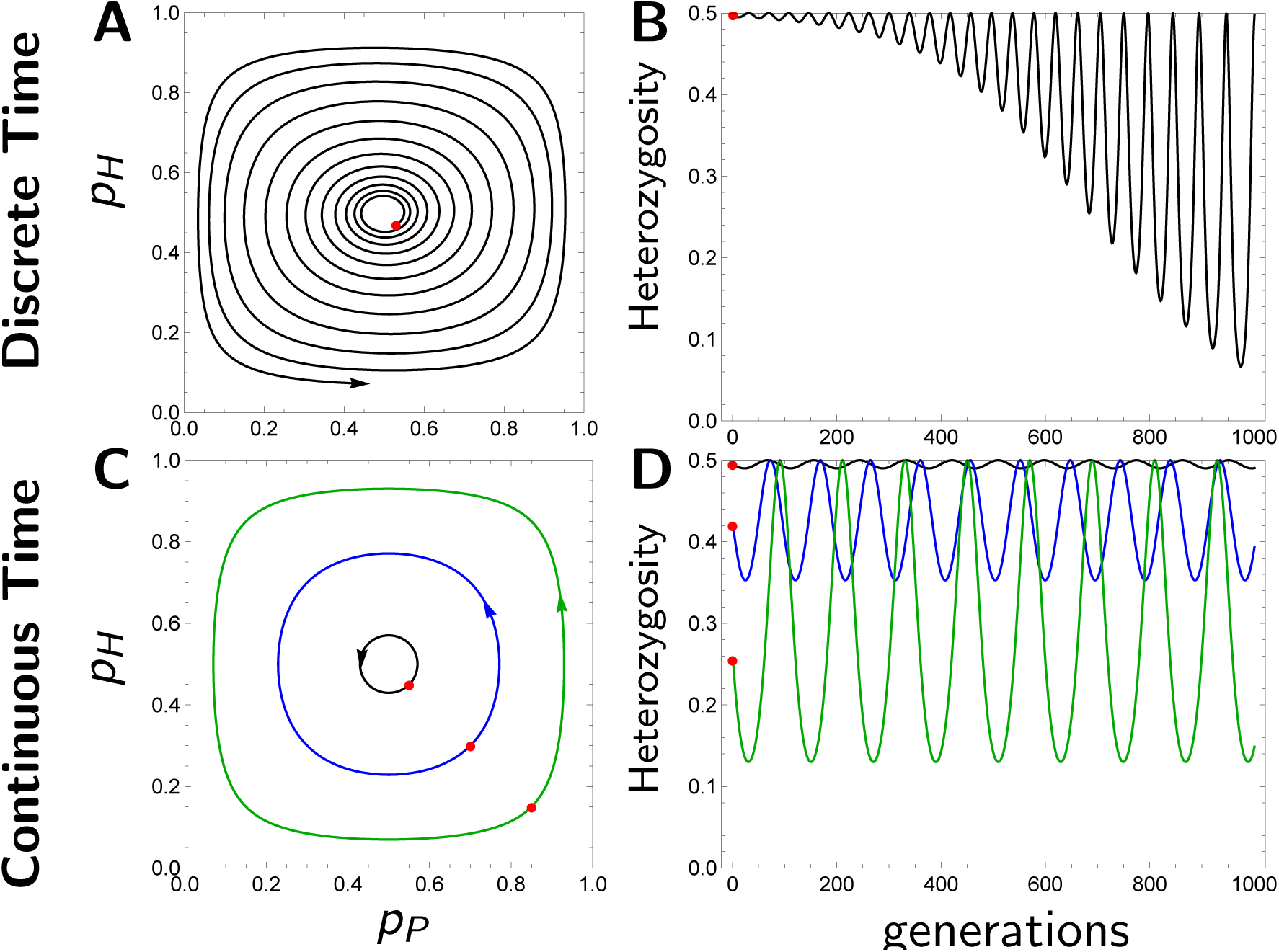
Deterministic dynamics of matching-alleles coevolution. Panel A: Phase-plane diagram of the unstable allele frequency cycles of the discrete-time matching-alleles model given by system (5) starting from a small initial perturbation (red point) from the polymorphic equilibrium. Parameters: *X* = 0.8, *Y* = 0.4, *α* = 0.4, *t*_*max*_ = 1000. Initial conditions: Black *p*_*H*_ (0) = 0.54, *p*_*P*_ (0) = 0.47. Panel B: Corresponding temporal dynamics of host heterozygosity. Panel C: Phase plane of neutrally stable allele frequency cycles arising in the continuous-time MAM given by system (3) starting form three different initial conditions shown by the red points. Parameters: *X* = 0.8, *Y* = 0.4, *κ* = 100, *δ* = 1, *α* = 0.2, *γ* = 1, *t*_*max*_ = 1000. Initial conditions: Black/Solid *p*_*H*_ (0) = 0.55, *p*_*P*_ (0) = 0.45, Blue/Dashed *p*_*H*_ (0) = 0.7, *p*_*P*_ (0) = 0.3, Green/Dotted *p*_*H*_ (0) = 0.85, *p*_*P*_ (0) = 0.15. Panel D: Corresponding temporal dynamics of host heterozygosity.

In contrast to the unstable Red Queen cycles generated in discrete time, analogous continuous-time models create neutrally stable allele frequency cycles (Woolhouse et al., 2002). Perturbations from the polymorphic equilibrium in these models lead to allele frequency cycles of constant amplitude, often referred to as a “dynamic polymorphism” (Hamilton, 1993). This dynamic polymorphism gives rise to corresponding cycles in heterozygosity also of constant amplitude (Figure 1C). This neutral stability indicates that, when averaged over the cycle, genetic variation remains constant in an infinite population, much like the behaviour of a neutral locus in a single-species model. We thus hypothesize that drift in a finite population may have similar effects in the coevolutionary model as in the neutral single-species model. Our aim here is therefore to develop an appropriate continuous-time MAM of coevolution in a finite population of constant size that, like the standard continuous-time MAM, does not lead to an inherent decline in heterozygosity in the deterministic limit. This model allows us to quantify the loss of heterozygosity with drift in the coevolutionary MAM for comparison to the standard single-species neutral model.

All the models discussed thus far make the traditional assumption that both host and parasite densities are infinite and controlled by factors independent of the host-parasite interaction. This is the case, for example, when host and parasite population sizes are fixed by a hard carrying capacity that is very large. The numbers of hosts and parasites may, however, vary dramatically in response to coevolution (Papkou et al., 2016). In addition parasites that are transmitted directly between hosts may be subject to epidemiological dynamics. While we have reason to believe that SIR dynamics can stabilize allele frequency dynamics (MacPherson et al., 2018), our goal in this work is to focus solely on the effects of NFDS that arises from coevolution. Unlike previous analyses, our goal here is to examine the stochastic nature of coevolutionary dynamics, allowing us to quantify rates of loss of genetic variation. To do so, we focus on the simplest case of strict external control of population size in both hosts and parasites without demographic or epidemiological feedback.

## 3. The Model

We use a continuous-time birth-death model to describe the coevolutionary dynamics between a host and a free-living pathogen, as depicted in Figure 2. To keep the total host and pathogen population sizes constant at the same fixed value *κ*, we use a Moran model design with coupled birth-death events. Extension of the model to unequal host and pathogen population sizes is straightforward and the results do not differ qualitatively from those presented here (*pers. comm*.). Both host and pathogen are haploid, with coevolution depending on a single biallelic locus in each species. We represent the number of hosts and pathogens of each type by *H*_*i*_ and *P*_*j*_ where *i* ∈ {1, 2} and *j* ∈ {1, 2}. Hosts of type *i* come into contact with pathogens of type *j* at a density-independent rate 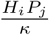. In keeping with MAM, upon contact the pathogen successfully infects the host with probability *β*_*i,j*_. If the host and pathogen carry “matching” alleles (*i* = *j*) then *β*_*i,j*_ = *X*, whereas “mis-matching” infection occurs with a reduced probability *β*_*i,j*_ = *Y < X* for *i /*= *j*. If infection occurs, hosts die instantaneously with probability *α*. The resulting rate of fatal infection of host type *i* from pathogen type *j* is 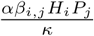. Fatal infections result in four coupled events 1) the death of the infected host of type *i*, 2) birth of a random host, 3) birth of the infecting pathogen of type *j* representing the release of infectious particles, and 4) death of a random pathogen. Non-fatal infections do not lead to the birth or death of either the host or pathogen, effectively assuming that the pathogen mounts a very limited infection and returns to the free-living pathogen population. In addition to these four events associated with infections, natural host death-birth and free-living pathogen death-birth occur at per-capita rates *δ* and *γ*, respectively. Both events are non-selective consisting of the death and birth of random host and pathogen genotypes.

**Figure 2:**
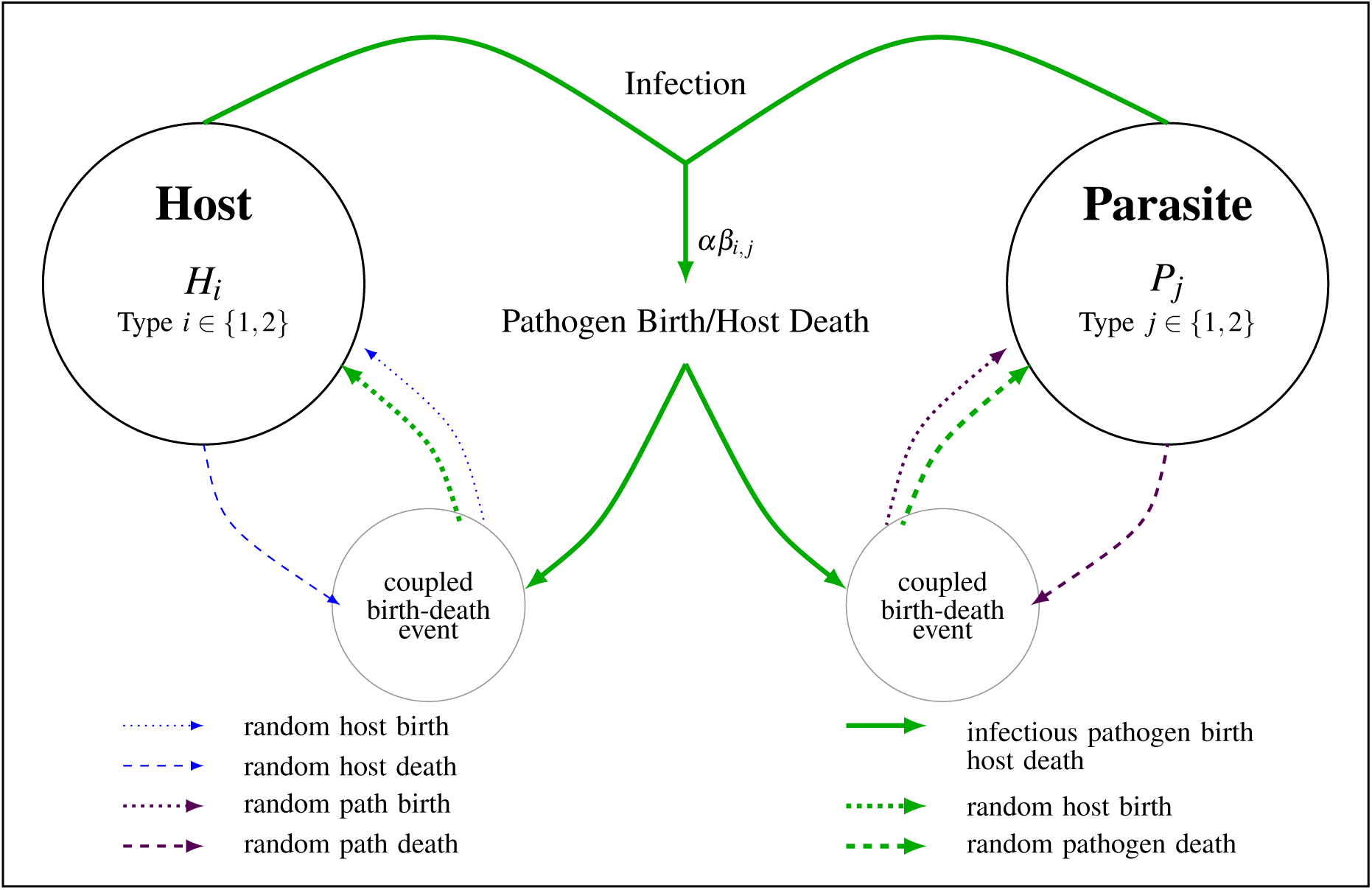
Schematic of the Moran MAM. Coevolution consisting of three different types of coupled birth-death events. Infection (green/thick), natural host death (blue/thin), and free-living pathogen death (purple/standard). Solid lines denote events that occur at a rate that depends on the genotype whereas dashed arrows represent birth and death of random individuals irrespective of their genotype.

As stated above, our primary aim is to compare the maintenance of genetic variation in a finite coevolving population to that expected under neutral genetic drift alone. Focusing on genetic variation in the host, for a population of finite size *κ*, the expected decline in host-heterozygosity from neutral genetic drift in the haploid Moran model is given by:

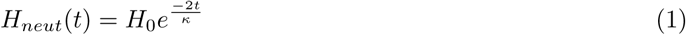

where *H*_0_ is the initial heterozygosity and time, *t*, is measured in units of host generations, (i.e. the expected time to *κ* deaths, *κ/*(*κd*) = 1*/d* where *d* is the death rate). To compare the neutral single-species model to a host-parasite model, we need to define time in the host-parasite model in an equivalent way. In the coevolutionary model, the per capita host death rate, *D*, hence the host generation time, changes over time as the allele frequencies change.

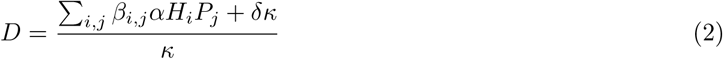

To measure time in terms of host generations we therefore rescale absolute time *τ* such that *t* = *Dτ* where *t* is again in units of host generations.

### 3.1. Deterministic dynamics

In the limit as the total host and pathogen population size goes to infinity (*κ → ∞*) the dynamics of the host and pathogen allele frequencies *p*_*H*_ = *H*_1_*/κ* (1 − *p*_*H*_ = *H*_2_*/κ*) and *p*_*P*_ = *P*_1_*/κ* (1 − *p*_*P*_ = *P*_2_*/κ*) are given by the following system of differential equations:

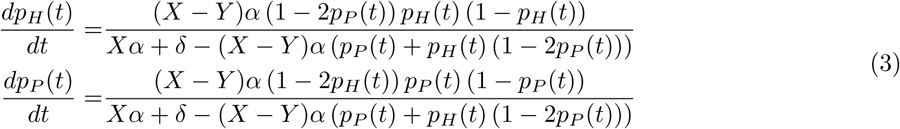

As shown in Figure 1, the behaviour of this Moran MAM in the deterministic limit is identical to that of the traditional continuous-time MAM (Woolhouse et al., 2002). Specifically, there are five equilibria of system (3). Four are unstable equilibria characterized by the fixation of one host and one pathogen genotype. The final polymorphic equilibria at 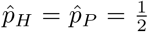 is neutrally stable with purely imaginary leading eigenvalues, generating neutral-limit cycles in allele frequencies (see Figure 1C).

As discussed previously, the dynamic polymorphism that arises in the deterministic continuous-time MAM neither maintains nor depletes host heterozygosity (see Figure 1D). While the neutral stability of the MAM is well known, the consequences on heterozygosity are under-appreciated. Contrary to the proposed effect of coevolution and NFDS on genetic diversity (Haldane, 1949; Clarke, 1979), neutrally stable allele frequency cycles neither deplete nor restore genetic variation. Rather, in an infinite population, coevolution has no net effect on genetic diversity, averaged across a cycle. To see if this behaviour holds in finite coevolving populations, we begin by describing an analytical approach to compare the dynamics of genetic diversity in a finite population to those expected under neutral genetic drift. Because this analytical approach only applies while the cycles are small we couple this approach with an individual-based simulation describing the loss of genetic diversity more generally.

### 3.2. Ensemble moment dynamics

To model the dynamics of genetic diversity in a finite population we express the model depicted in Figure 2 as a continuous-time Markov chain. The complexity of the model limits the available analytical approaches. One approach we can use, however, is the ensemble moment approximation, the derivation of which is given in the Appendix. The result is an approximation for the expected host heterozygosity which we will call 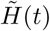. As with the neutral expectation in equation (1), this is the expectation of host heterozygosity over all realizations of the stochastic process, the approximation of which is given by a Taylor series assuming the system is near the deterministic equilibrium (*H*_0_ ≈ 1*/*2) and the population size large.

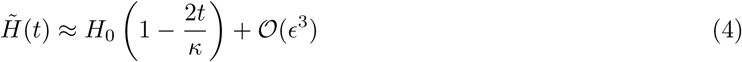

where *ϵ* is the deviation of the stochastic process from the deterministic equilibrium and *ϵ* = 1*/κ*. Once again *t* is measured in units of host generations. To this order, the ensemble moment dynamics are identical to that of the neutral expectation given by (1). Hence, as expected from the deterministic dynamics, host heterozygosity in the MAM declines as expected under neutral genetic drift. Although analytically unwieldy, numerical evaluation of 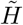 to third order reveals slight deviations between the ensemble moment dynamics and the neutral expectation that are also observed in the individual-based simulations presented in the following section. Specifically, 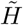 exceeds the neutral expectation when the free-living pathogen death rate *γ* is less than the death rate of the host *δ* (negative values of *γ* − *δ* in Figure 3B) and falls below the neutral expectation as this rate increases.

**Figure 3:**
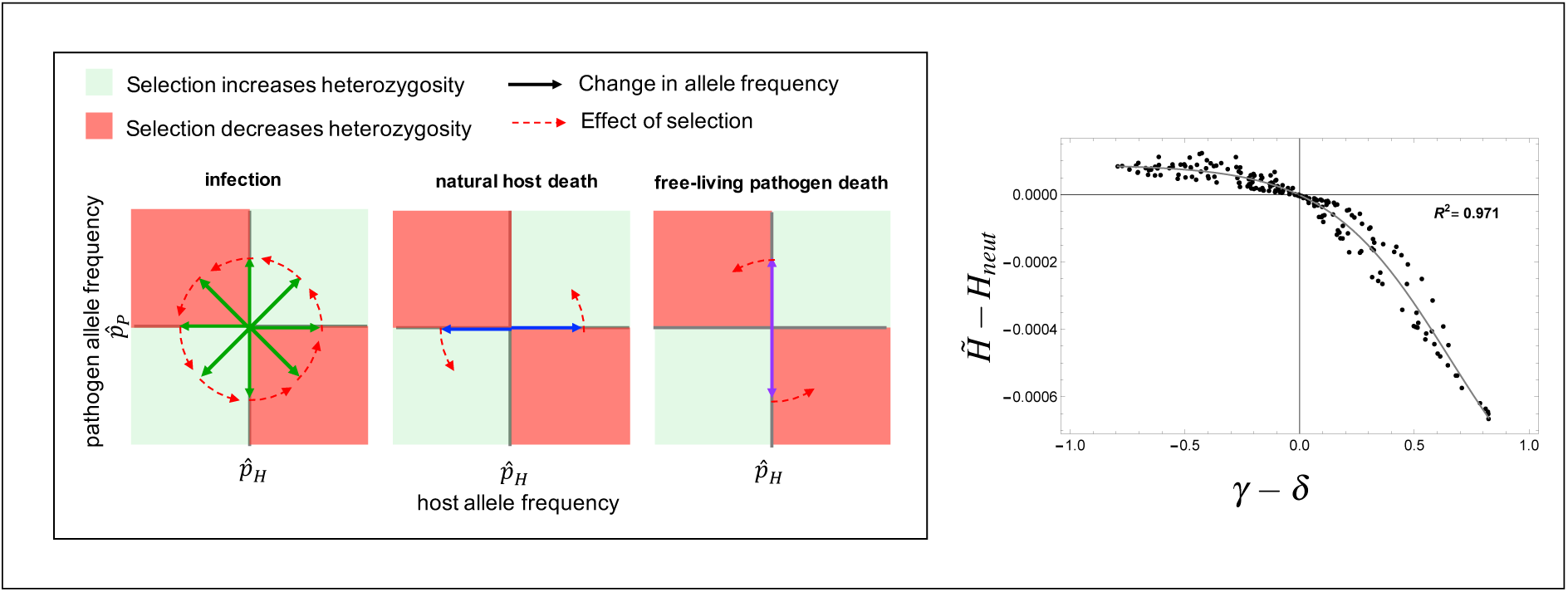
The effect of early perturbations. Shown schematically in the left panel are the effects of the three processes, infection, natural host death, and natural pathogen death on host and pathogen allele frequency and the resulting effect of selection on host heterozygosity. 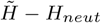 as a function of the relative host and pathogen death rates is shown in the right panel. Points give the deviation at time *t* = 50 host generations for 200 randomly drawn combinations of *γ* and *δ*, where 0.1 *< γ <* 1, 0.1 *< δ <* 1, and *X* = 0.9, *Y* = 0.7, *α* = 0.2, *κ* = 10^5^.

The mechanism behind these lower-order deviations is shown schematically in Figure 3A. If we start, for example, at the equilibrium host and pathogen allele frequencies 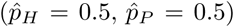, infection, natural host death-birth, and free-living pathogen death-birth events perturb the allele frequencies. Selection then acts on those perturbations to produce small counter-clock wise allele frequency cycles. Although over the long term these allele frequency cycles have no net effect on heterozygosity, they do have an effect over the short term. For example, if only the host allele frequency is perturbed by a random host birth event then during the first quarter cycle natural selection will increase host heterozygosity, because the parasite that matches the host that, by chance, became more common, will spread in turn reducing the more common host genotype. In contrast, if only the pathogen allele frequency is perturbed, selection will decrease host heterozygosity by preferentially killing off the host that matches the parasite allele that increased by drift.

The effect of these early quarter cycle responses to selection would average out if perturbations were distributed uniformly about the equilibrium. However, natural host death affects only the host allele frequency whereas free-living pathogen death only affects the pathogen allele frequency. Hence the relative rates of these two events determine whether these early responses to selection transiently increases or decreases heterozygosity. Specifically, relatively high rates of natural host death and low rates of free-living pathogen death (*γ* − *δ <* 0) lead to an increase in host heterozygosity relative to the neutral expectation (see Figure 3B).

### 3.3. Individual-based simulations

Using a Gillespie algorithm we simulated host-parasite coevolution in a finite population. For each of one-hundred randomly drawn parameter combinations, we simulated coevolution for seven different populations sizes ranging on a log scale from *κ* = 10^2^ to *κ* = 10^3.5^ = 3162. The relative infection rates of matching and mis-matching genotypes were drawn such that 0 *≤ Y < X ≤* 1. We restricted the probability of dying from infection, *α*, to lie between 0 and 1*/*2 as much stronger selection causes the rapid loss of alleles, limiting the amount of coevolution. Host and pathogen free-living natural death rates *δ* and *γ* were both drawn between 0.1 and 1. For each of the 700 parameter combinations, we simulated 1000 replicate populations for *t*_*max*_ = 500 host generations.

In tandem with each coevolutionary replicate population we simulated an analogous “neutral population” to check that it behaved correctly according to the neutral expectation given in equation (1). In these simulations, the exact same series of host and pathogen birth-death events occurred as in the coevolutionary simulations, except that the individual who was born or died due to infection was drawn at random, these simulations captured random genetic drift in both host and pathogen. As shown in Figure 4B the mean heterozygosity of these neutral simulations (shown in gray) are identical to the analytical neutral expectation given by equation (1), confirming that this is the appropriate neutral expectation for the two-species model.

**Figure 4:**
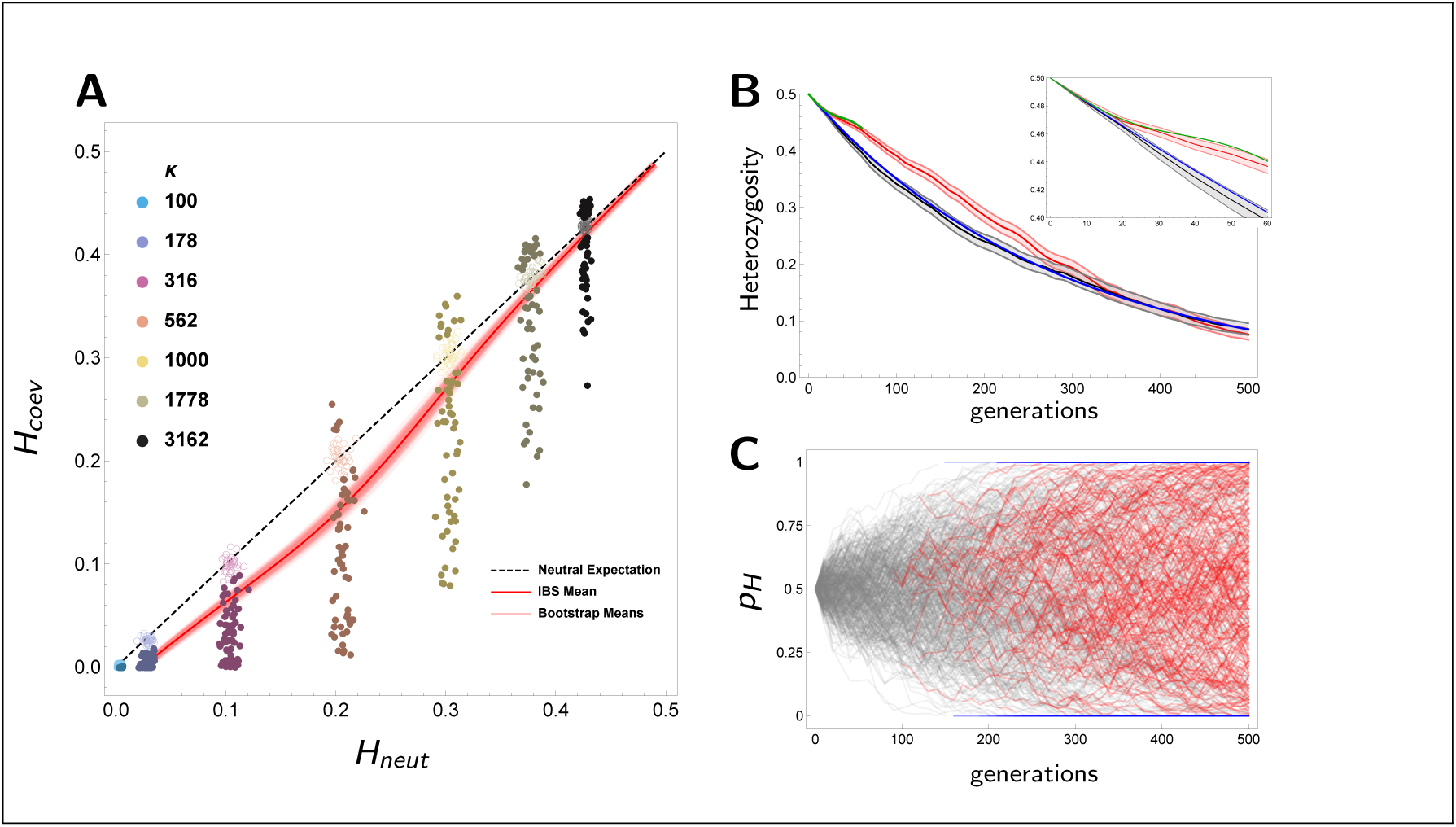
Individual-based simulations. Panel A: Heterozygosity in the coevolving populations versus the neutral expectation at time *t* = 250 host generations. Black dashed line represents the expectation under neutrality. The red curve is the smoothed fit of the mean, light red lines are bootstrap means. The points are the mean heterozygosities for each of the 420 parameter combinations averaged across the 1000 replicate populations simulated for each. Points are filled if the mean heterozygosity for the parameter combination differs significantly from the neutral expectation (with 95% confidence with Bonferroni correction for the 100 points for each *κ*) versus empty when the replicate population means overlap neutrality. Panel B: Example dynamics for a single parameter condition (*X* = 0.29, *Y* = 0.14, *α* = 0.38, *δ* = 0.99, *γ* = 0.24, and *κ* = 10^2.75^ = 562). Red region represents CI of heterozygosity in the coevolving population, gray region is the CI in the analogous neutral simulation, Blue line is the neutral expectation from equation (1) and green line is heterozygosity in the second order ensemble moment approximation. Panel C: Allele frequency dynamics for example simulation. Gray trajectories represent ongoing coevolution, red are periods of directional selection, while blue are fixed host replicate populations. Parameters: *X* = 0.26, *Y* = 0.25, *α* = 0.35, *δ* = 0.36, *γ* = 0.83, *κ* = 10^3.5^ = 1778

Figure 4 captures the four key results of our model. First, as predicted by the second-order ensemble moment approximation given in equation (4) and shown in the inset of panel B the dynamics of heterozygosity in the coevolving populations behave neutrally, for approximately the first 10 generations, when the dynamics are very near the deterministic equilibrium. Second, as the distance from the deterministic equilibrium grows, the coevolutionary dynamics depart slightly from the neutral expectation due to the transient effects of selection on perturbations. The resulting slight increase (decrease) in heterozygosity is shown by the filled points in panel A that are either slightly greater (less) than the neutral expectation. The third key result is that heterozygosity is sometimes substantially less than expected from the neutral expectation. Following fixation, the matching-alleles genetic mechanism becomes one of directional selection, favouring the mismatching host of the remaining pathogen genotype. This directional selection (shown by red trajectories in Figure 4C) in turn rapidly erodes genetic variation within the host population. Finally, as predicted by the deterministic model, host heterozygosity in most cases is near the neutral expectation. Specifically, open points in Figure 4A) indicate parameter conditions that did not differ significantly from the neutral expectation.

## 4. Discussion

Contrary to theories that posit that host-parasite coevolution and the associated negative frequency-dependent selection (NFDS) should maintain genetic variation (Haldane, 1949; Clarke, 1979; Takahata and Nei, 1990), we use stochastic methods, both describing ensemble moments and conducting individual-based simulations, to show that genetic variation is lost at nearly the same rate as in the neutral model (continuous time) or faster (discrete time). Although these results are consistent with stability analyses of the corresponding deterministic models, such stability analyses only apply near the equilibrium with equal allele frequencies. Thus, in contrast to single-species NFDS, coevolutionary NFDS does not generally maintain genetic variation. Specifically, the neutrality of the cycles ensures that for any given host allele frequency selection is as likely to be selecting for as against that allele. While the largely neutral effect of matching-alleles coevolution is recovered in our models of finite-population size, we find that stochasticity introduces two new effects. First is the effect of the initial response to selection following perturbations in allele frequency from the deterministic equilibrium, which may on average either maintain or deplete heterozygosity depending on the relative death rates of the host versus the pathogen. The second non-neutral effect is that of directional selection following pathogen fixation which rapidly erodes genetic variation.

The contrasting effects of coevolutionary indirect-NFDS and single-species direct-NFDS have been pointed out previously by Brown and Tellier (2011) in the context of the gene-for-gene coevolutionary model. Nevertheless, the distinction between these two processes and in particular their differing effects on the maintenance of genetic variation remains under-appreciated. Our finding that the MAM does not inherently maintain genetic variation contrasts with and clarifies the findings of previous computational models of the maintenance of genetic variation at MHC loci (Ejsmond and Radwan, 2015; Borghans et al., 2004). Simulating matching-alleles coevolution at multiple loci in the presence of rapid host mutation, both Ejsmond and Radwan (2015) and Borghans et al. (2004) find a weak signal of increased fitness of rare host alleles. Our results clarify, however, that this is not in fact a true signal of negative frequency-dependent selection. Rather this is a result of the fact that novel, and hence rare, host alleles may be favoured transiently if there does not yet exist genetic variation at the corresponding pathogen locus. In contrast, long term changes in host allele frequencies are not associated with corresponding changes in fitness as would be required under NFDS (see Borghans et al. (2004) Fig 4A).

Here we have focused on coevolution in a single population. Coevolution is however inherently a spatial process. Previous theoretical models suggest that spatial structure in combination with variability in the strength and nature of host-parasite coevolution across space may promote local adaptation (Gandon et al., 1996; Nuismer, 2006). Our results illustrate, however, that this is not the result of coevolution within each individual deme but rather an emergent effect of spatial structure and gene flow. Whether these spatial models maintain more genetic variation than expected under neutral drift remains a valuable topic for future work.

In the main text we have focused on host-parasite coevolution in a continuous-time model as opposed to a model of coevolution in discrete time (see equation A9). This is because, as explored in the background section, the continuous-time model is more likely to maintain genetic variation based on the behaviour of the deterministic dynamics. We confirmed that this expectation holds when extended to finite populations by simulating host-parasite coevolution in a Wright-Fisher MAM (see Appendix). As expected host-heterozygosity in such a discrete-time MAM is significantly less than the neutral expectation and less than observed in the continuous-time model (see Figure 5).

**Figure 5:**
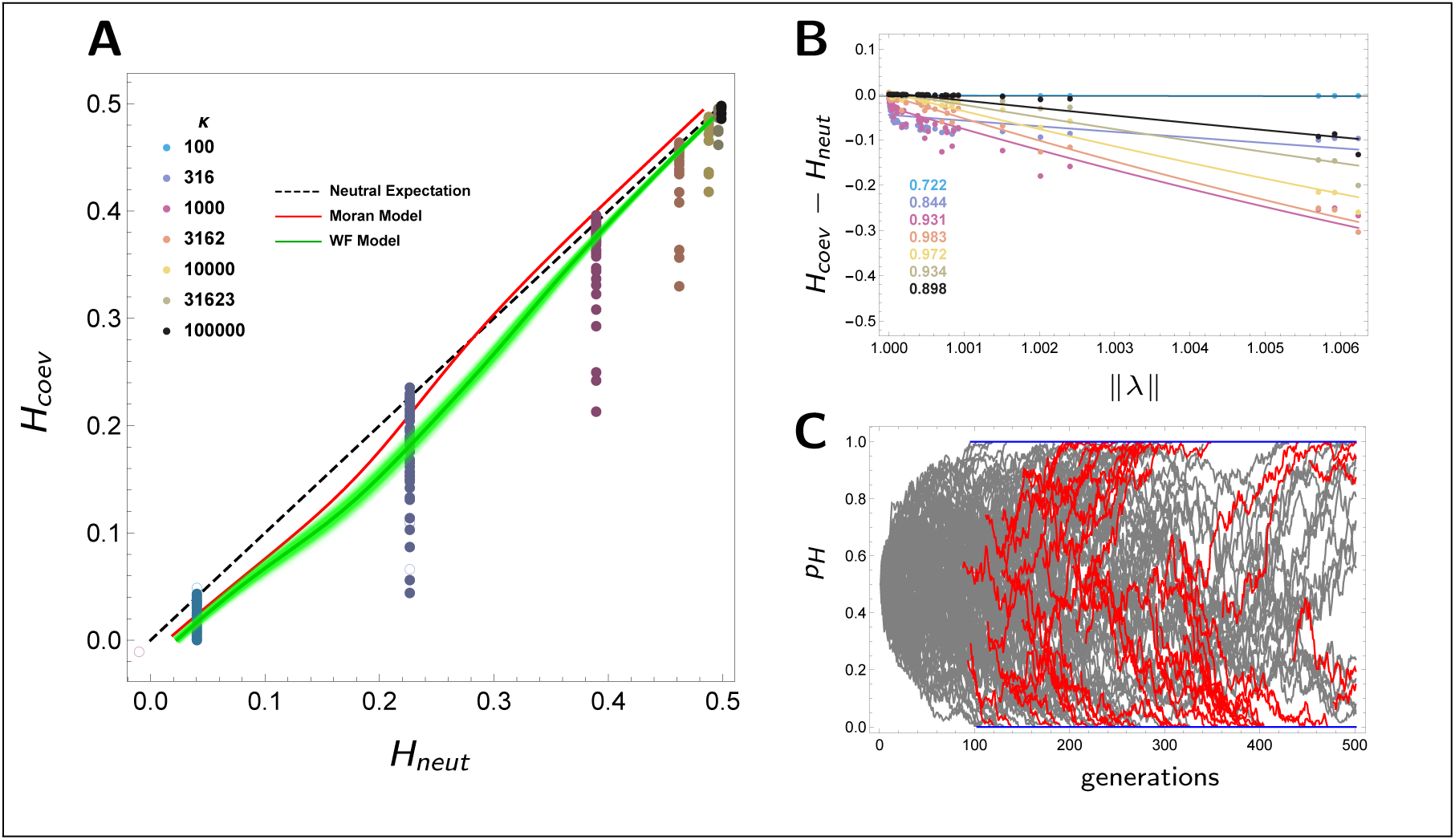
Comparison between discrete- and continuous-time models. Panel A: Mean heterozygosity in a coevolving population (at *t* = 250 generations) versus the neutral expectation for the Moran model in continuous time (red as in- see Figure 4) versus the Wright-Fisher model in discrete time (green), light green lines give smoothed fits to 1000 bootstrap samples. Points depict heterozygosity for each of the 420 parameter conditions simulated for the WF model averaged over the 1000 replicate populations. Panel B: Deviation in heterozygosity from the neutral expectation in the WF as a function of the magnitude of the leading eigenvalue. *R*^2^ values are given for the non-linear model fit of *a* + *e*^−*b*(*x*−1)^. Panel C: Example allele frequency dynamics in the WF model. Allele frequency trajectories are coloured gray when there is ongoing coevolution, red when there is directional selection, and blue when the host population is fixed. Parameters: *X* = 0.9, *Y* = 0.26, *α* = 0.047, *κ* = 10^2.5^ = 316

The largely neutral effects of the continuous-time MAM and variation eroding effects in discrete-time makes coevolutionary NFDS an insufficient explanation for the high levels of genetic diversity at host immuno-defence loci. The models presented here make several implicit assumptions that may influence the maintenance of genetic diversity. In particular, we assume that the population size remains constant. Although this is a classic assumption in coevolutionary theory, coevolution is known to have drastic effects on population sizes of both host and pathogen in natural systems (Papkou et al., 2016). Indeed inclusion of explicit population size dynamics in the MAM results in fluctuations in host and pathogen population size (Nuismer, 2017; Frank, 1993). While these changes in population size are expected to have little to no effect on the allele frequency dynamics, they are expected to have important consequences on the maintenance of genetic variation (Crow, 1970). We also assume that the host and parasite generations are synchronized. The extensive literature on the deterministic stability of the polymorphic equilibrium has suggested that temporal asynchrony between the host and parasite, arising for example through seed dormancy, can stabilize polymorphisms and maintain variation (Brown and Tellier, 2011; Tellier and Brown, 2009). Developing an analogous MAM model, we find that here too including a temporal delay in the ability of one species to respond to the other (here by including seed dormancy) has a stabilizing effect and can help maintain variation (see Appendix).

The maintenance of genetic diversity at human MHC loci may also be the result of coevolution between hosts and pathogens that are transmitted directly from one host to another and hence subject to epidemiological dynamics. Our previous work showed that these epidemiological dynamics stabilizes Red Queen allele frequency cycles resulting in a stable polymorphic equilibrium (MacPherson et al., 2018). In contrast to the instability of the discrete-time MAM and the neutral stability of the continuous-time MAM, the stabilizing effects of these epidemiological dynamics are expected to maintain genetic variation. While the work presented here was aimed solely at understanding the effects of coevolution on the maintenance of genetic variation in finite populations, extending these results to include more realistic population size and epidemiological dynamics is necessary to fully understand the maintenance of genetic variation in natural systems.

## Acknowledgements

We thank Matt Pennell, Jonathan Davies, Jeff Joy, Dan Coombs, and Aurélien Tellier for their many helpful suggestions that improved this manuscript. This project was supported by a fellowship from the Godfrey-Hewitt Mobility Award and the AAUW dissertation fellowship and a fellowship from the University of British Columbia to A.M. as well as a Natural Sciences and Engineering Research Council of Canada grant to S.P.O. (RGPIN-2016-03711).

### Appendix

#### A1. The Ensemble Moment Approximation

##### A1.1. General Approach

We consider the general stochastic process in *Z*^+*n*^. Specifically, for the MAM model presented here *n* = 2, one dimension for the number of hosts of type 1 and pathogens of type 1, respectively. Let 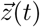 denote the state of the process at time *t*, for our case 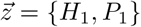. We begin by transforming the variables into their deviation from the deterministic equilibrium. Specifically if 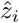 is the value of variable *i* at equilibrium, then the transformed variables are given by 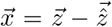. In addition, we only consider the case where the initial condition of the stochastic processes is the deterministic equilibrium, 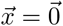. This transformation in variables will then allow us to easily express the moments of the stochastic process in terms of Taylor series around the deterministic equilibrium. In terms of the transformed variables, the stochastic process is described by *m* distinct events that occur at rates 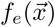 where *e* = {1, 2, …, *m*}. Finally, we denote the ensemble average, the average across all sample paths, by 〈*〉.

The aim of the ensemble moment approximation is to derive a system of ODEs giving the change in the moments of the stochastic process. Beginning with the first moment (mean) we have:

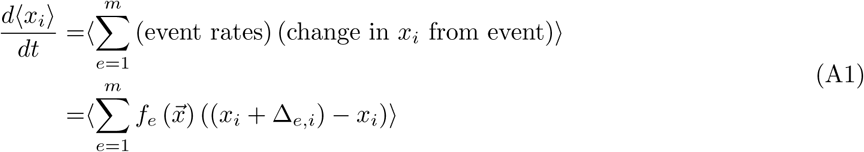

where Δ_*e,i*_ is the change in *x*_*i*_ from event *e* (Keeling, 2000). To take the ensemble average of the sum we express the argument as a polynomial by approximating it with a Taylor series around the deterministic equilibrium assuming *x*_*i*_ ∀ *i* is small. Equation (A1) is then approximated as:

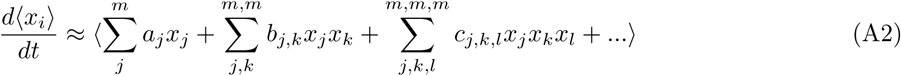

where *a*_*j*_, *b*_*j,k*_, and *c*_*j,k,l*_ are constants, which are themselves functions of the deterministic equilibrium 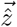. Rearranging we have:

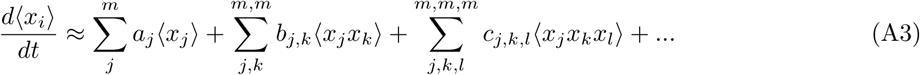

The result is an expression for the change in the first moment as a linear function of the first, second, third, and higher moments.

We use the same technique to derive the second moment ODEs:

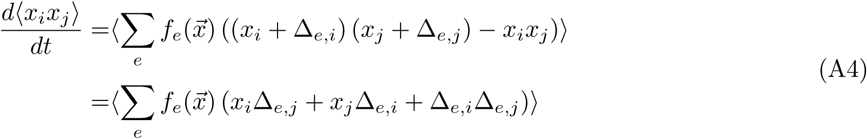

Approximating the right-hand argument as a sum once again by a Taylor series expansion we have:

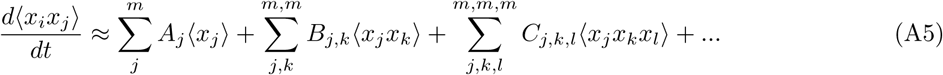

Similarly for the third moments:

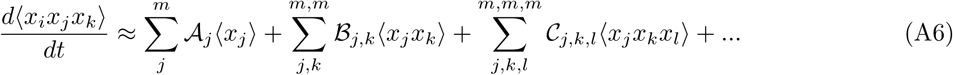

where *A, B, C, 𝒜, B*, and *𝒞* are all constants that depend on the deterministic equilibrium.

##### A1.2. Dynamics of host heterozygosity

We are interested in the dynamics of host heterozygosity and calculate the ensemble moment approximation for the dynamics of heterozygosity, 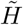:

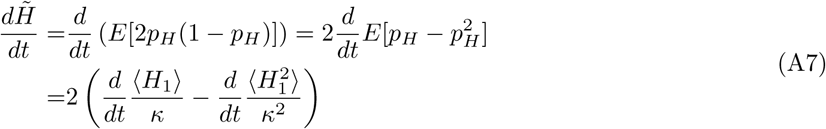

The dynamics of heterozygosity thus depends directly on that of the first two moments. If we substitute in equation (A3) and (A5) and assume that population size is large *κ* = 1*/∈*, then approximated to *mathcalO*(*∈*^2^) the dynamics of host heterozygosity simplify to equation (4) given in the main text.

#### A2. Maintenance of Genetic Variation in Discrete Time

##### A2.1. Model specification

We model host-parasite coevolution in discrete time with a Wright-Fisher model with a constant host and pathogen population size *κ*. In each generation, hosts come into contact with a single random pathogen. If host and pathogen have the same genotype *i*, the probability of successful infection is given by *β*_*i,i*_ = *X*, whereas hosts of type *i* are infected by pathogens of mis-matching genotype *j*, with probability *β*_*i,j*_ = *Y < X*. In the absence of infections host’s and pathogen’s have a fitness 1, whereas infection decreases host fitness by a factor *α*_*H*_ and increases pathogen fitness by *α*_*P*_ such that the expected fitness of host genotype *i* and pathogen genotype *j* are given by:

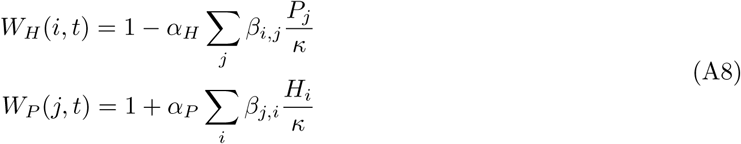

##### A2.2. Deterministic dynamics

As we do for the continuous-time model, we begin by developing an understanding of the effect of coevolution on genetic variation in a finite population by analysing the dynamics of heterozygosity in the limit as the population size goes to infinity. In this deterministic case the coevolutionary dynamics are given by the following difference equations:

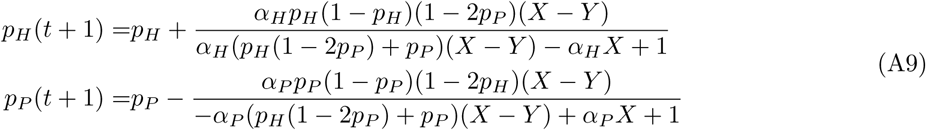

There are five equilibria of this model, four of these specify the fixation/loss of one host and one pathogen type, and a fifth internal equilibrium occurs at *p*_*H*_ = *p*_*P*_ = 1*/*2. Unlike the continuous time model explored in the main text this internal equilibrium is unstable and cyclic. The dynamics of the allele frequency and heterozygosity are shown in Figure 1 panels A and B, respectively. The rate at which the amplitude of the cycles grow is given by the magnitude of the leading eigenvalue, which is given by ∥*λ*∥, a result consistent with those of (M’Gonigle et al., 2009).

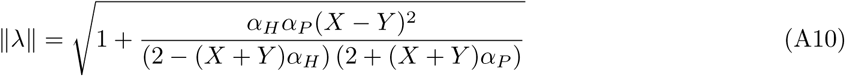

Due to the instability of the internal equilibrium host heterozygosity decays over time. Furthermore the larger the value of ∥*λ*∥ the faster the heterozygosity should decay.

##### A2.3. Simulations of a finite population

In analogy to the continuous-time model presented in the main-text where the death of the host and birth of the pathogen from infection are coupled, for our simulations in a finite population we assume *α*_*H*_ = *α*_*P*_ = *α*. As with the continuous-time model we simulate coevolution for 100 random parameter combinations of *X, Y*, and, *α* (0 *< α <* 0.5, 0 *< Y < X <* 1) and 7 values of *κ* (ranging on a log scale between 10^2^ and 10^5^) for a total of 700 simulations. In contrast to the continuous-time model natural host death and free-living pathogen birth are incorporated directly in the Wright-Fisher model. For each of the simulations, sample paths were generated for 1000 independent replicate populations. All simulations are initialized at the internal polymorphic equilibrium and run for *t* = 500 generations.

In the absence of coevolution, the dynamics of host heterozygosity are given by the neutral expectation for the Wright-Fisher model:

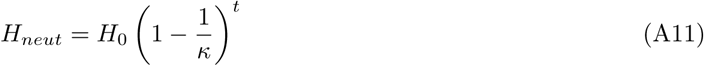

The difference between the simulated heterozygosity and the neutral expectation at time *t* = 500 is shown in Figure 5A. When population sizes is large (*κ* = 10^5^ upper right hand corner) the deviation in heterozygosity between the coevolutionary simulations and neutral expectation, Δ*H* = *H*_*coev*_ − *H*_*neut*_, is statistically negative as expected due to the erosion of genetic variation by coevolution. The magnitude of this deviation is however small, so while coevolution does indeed erode genetic variation the process is slow with only small effects over the parameter and time scale sampled.

As with the continuous-time model presented in the main text, directional selection (see Figure 5C) following fixation in the pathogen has a much larger effect than that of coevolutionary selection. When the population size is small (e.g. *κ* = 10^2^), drift dominates over the effects of selection, and the heterozygosity behaves nearly neutrally. As the population size increases, the effect of this directional selection emerges. As predicted by the deterministic model variation in Δ*H* for a given population size is explained largely by variation in ∥*λ*∥ (see Figure 5B). This is not only because ∥*λ*∥ determines the effect of coevolution but \because it is highly correlated with the probability of pathogen fixation and the strength of directional selection following pathogen fixation.

#### A3. Seed Dormancy

Here we extend the above discrete-time model to include the effects of seed dormancy, or any other process that causes a subset, *s*, of the host population to avoid parasite induced selection, making the host less responsive during coevolution. We analyse the effect of the resulting temporal asynchrony between host and parasite on the stability of the polymorphic equilibrium. Given a life cycle shown if Figure 6 we model evolution by following the allele frequencies among the host seedlings *p*_*H*_, host juvenile seeds *p*_*J*_, and within the parasite *p*_*P*_. The dynamics are given by:

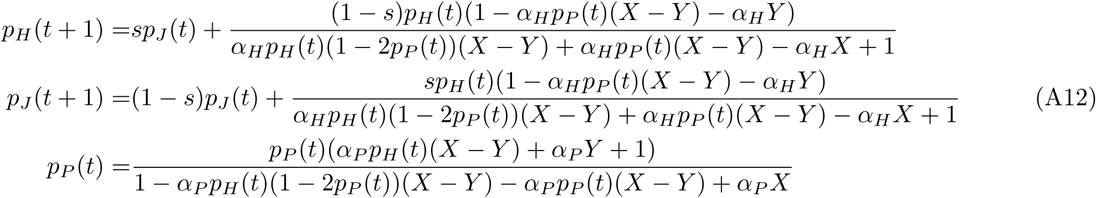

**Figure 6:**
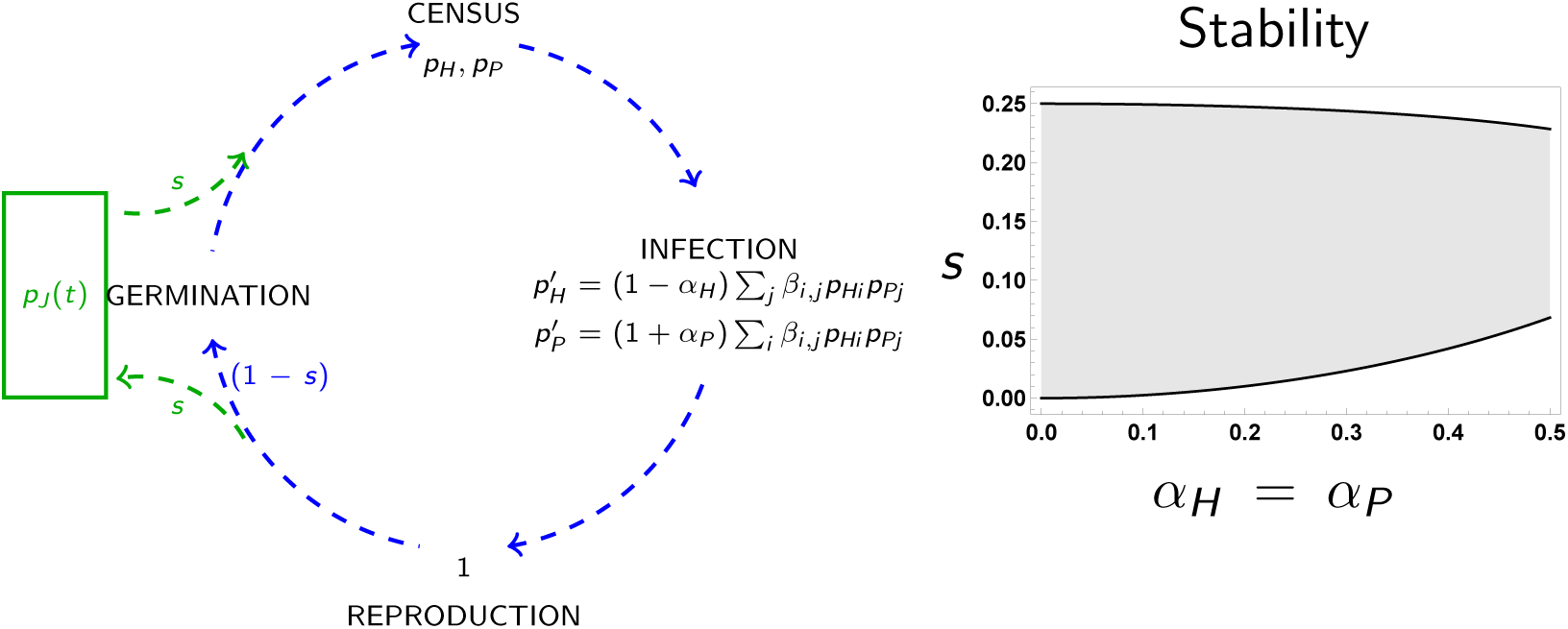
Life-cycle diagram and numerical stability analysis of MAM with seed dormancy. Host and pathogen life-cycle (blue arrows), flux in and out of seed bank (green arrows). Shaded region of right-hand plot gives region for which the polymorphic equilibrium is stable assuming *α*_*H*_ = *α*_*P*_ and given *X* = 1, *Y* = 0.

There are five equilibria of system A12, four of which are trivial fixation/loss of the host and pathogen. The fifth internal equilibrium is at 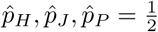. The Jacobian matrix at this internal equilibrium is given by:

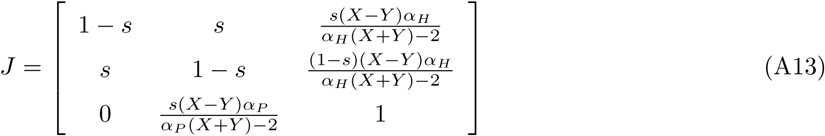

Focusing on the diagonal elements, we note that 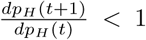 and 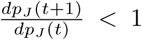. As noted by Tellier and Brown (2009) this is indicative of direct negative frequency-dependent selection. We can evaluate the stability of system A12 by using the Routh-Hurwitz conditions on the transformed characteristic polynomial of A13. Specifically, although the Routh-Hurwitz conditions can only be applied to identify the stability criteria in continuous time, if we transform the characteristic polynomial, *p*(*λ*), into a third order polynomial in *z* such that:

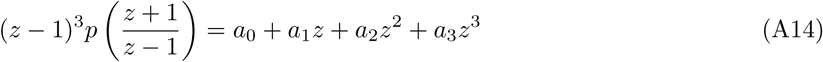

then the equilibrium is stable (all eigenvalues have a magnitude less than 1) if and only if *a*_*i*_ *<* 0 ∀*i* and *a*_2_*a*_1_ − *a*_3_*a*_0_ *>* 0 assuming *α*_*H*_ = *α*_*P*_ = *α*. These criteria are satisfied in our model for a range of *s*, indicating the the internal equilibrium is stable under these condition:

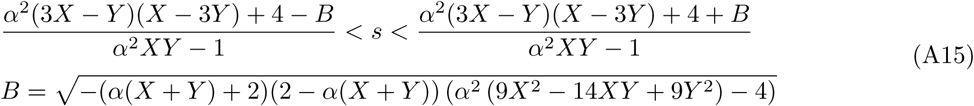

where *B* must be real.

#### A4. Supplementary Figures

